# Cardiac Atrial Compartmentalisation Proteomics: A Modified Density Gradient Method to Analyse Endo-lysosomal Proteins

**DOI:** 10.1101/2021.02.22.432193

**Authors:** Thamali Ayagama, Samuel J Bose, Rebecca A Capel, David A Priestman, Georgina Berridge, Roman Fisher, Antony Galione, Frances M Platt, Holger Kramer, Rebecca A B Burton

## Abstract

The importance of lysosomes in cardiac physiology and pathology are well established, and evidence for roles in calcium signalling are emerging. We describe a label-free proteomics method suitable for small cardiac tissue biopsies based on density-separated fractionation, which allows study of endo-lysosomal (EL) proteins.

Density gradient fractions corresponding to tissue lysate; sarcoplasmic reticulum (SR), mitochondria (Mito) (1.3 g/ml); and EL with negligible contamination from SR or Mito (1.04 g/ml), were analysed using Western Blot, enzyme activity assay and LC-MS/MS analysis (adapted discontinuous Percoll, and sucrose differential density gradient).

Kyoto Encyclopedia of Genes and Genomes, Reactome, Panther and Gene Ontology pathway analysis showed good coverage of RAB proteins and lysosomal cathepsins (including cardiac-specific cathepsin D) in the purified EL fraction. Significant EL proteins recovered included catalytic activity proteins. We thus present a comprehensive protocol and dataset of guinea-pig atrial EL organelle proteomics using techniques also applicable for non-cardiac tissue.

## Introduction

The concept of understanding proteins from a compartmentalisation perspective, their interconnected properties and dynamic distribution in health and disease is critical for deciphering the phenotype of a cell^1^. Significant advances in mass spectrometry-based proteomics allow scientists to achieve multidimensional measurements of proteins with greater efficiency, enabling for example the generation of more detailed maps of the human proteome^2^. Relative quantification methods of samples include label-free quantification^3^, *in vivo* metabolic stable-isotope labelling^4^, stable-isotope labelling using chemical tags that are covalently attached *in vitro*, tandem mass tags and isobaric tags for relative and absolute quantification (recently reviewed by Larance and Lamond, 2015^1^).

Methods available to analyse subcellular protein localisation in cells and tissues are diverse^5,6^. Depending on the cell or tissue to be analysed, the different methods have distinctive advantages and disadvantages. The subcellular fractionation methods most commonly combined with mass spectrometry-based analysis include differential centrifugation and either equilibrium gradient centrifugation or non-equilibrium gradient centrifugation^1^. One of the challenging issues encountered with subcellular fractionation is due to very small density differences between individual organelle fractions^7^. ^8^Advanced methods, such as localization of organelle proteins by isotope tagging (LOPIT^9^), offer advantages to differentiate large organelles, small intracellular vesicles and even large complexes such as ribosomes, purely based on their density and not requiring isolation and purification of organelles. An increased understanding of the physiological and structural interactions between intracellular organelles, such as the role of membrane contact sites (MCS)^8^ and inter-organelle nanojunctions in regulating physiological function^10^, also raises considerations for determining fraction purity. For example, Niemann-Pick type C protein 1 is now known to play a role in regulating MCS between lysosomes and the endoplasmic reticulum (ER)^8^. Such interactions raise the possibility for example of contamination of ER fractions by lysosomal proteins as a result of MCS formation.

Over recent years, significant progress has been made in establishing a region and cell-type resolved quantitative proteomic map of the human heart^11^. The value of such approaches has been demonstrated by the application of these data to define molecular changes in patients suffering from cardiovascular disease and to provide comparisons with known genomic parameters for cardiovascular disease including heart failure and atrial fibrillation (AF)^12,13^. Mishandling of Ca^2+^ regulation in cardiac cells is closely linked to the pathophysiology of cardiac arrhythmias such as AF^14^ and there is increasing evidence for an involvement of lysosomes in cardiac Ca^2+^ handling/mishandling^15,16^. In order to understand the contribution of the organelles involved in Ca^2+^ regulation (including lysosomes in addition to SR and mitochondria) of atrial function, a more detailed organelle-specific approach is required. The guinea pig (*Cavia porcellus*) is a common and well appreciated small mammal model used in cardiovascular research and recent work from our own group has highlighted the value of using *C. porcellus* tissue to study atrial Ca^2+^ handling in atrial cells^14,17^. *C. porcellus* cardiomyocyte electrophysiology, which includes the typical long plateau phase of the action potential is closer to that of human compared to mouse or rat. Our proof-of-concept study offers the advantage of scalability, involving utilisation of very small quantities of heart biopsies, for instance those obtainable during surgery. Using this approach, we will be able to explore the contribution of lysosomes in health and disease. In doing so we hope to unravel novel mechanistic insights, relating to changes in protein composition in diseased models that can then be related back to functional data.

The importance of lysosomes in cardiac physiology, both in health and in disease has long been recognised^18^. Early elegant ultrastructural and biochemical studies investigated the levels of lysosomal enzyme activity in many organs including the heart^19–22^. As early as 1964 an increased number of lysosomes was observed in the atrial muscle of chronically diseased or stressed hearts with acquired heart disease such as mitral stenosis^23^. In addition, Kottmeier *et al.* conducted studies in a dog model of atrial septal defects and their data demonstrated an increase in the number of myocardial lysosomes in cells subjected to increased metabolic demands^24^. The correlation however between the degree of stress and elevation in lysosome count could not be determined from these early studies. In 1977, Wildenthal^25,26^ looked at differences in cardiac lysosomal enzymes in detail and confirmed previous observations^19,27^ correlating increased age with the total activity of the lysosomal proteinase, cathepsin D, further highlighting links between lysosomal function and cardiac disease.

Development of techniques that facilitate proteomic characterization of individual organelles^28^ could provide valuable information regarding the function of lysosomal pathways in normal and disease states^16^. For instance, lysosomal calcium signalling via the nicotinic acid adenine dinucleotide phosphate (NAADP) pathway^29,30^.

Relatively little is known about the protein composition of the lysosomes in cardiac atria. In this study we developed a flexible, low-cost, modified density gradient method for endo-lysosomal organelle isolation, allowing better organelle protein identification from the processing of small amounts of frozen cardiac atrial tissue biopsies. We performed label-free, quantitative mass spectrometry that allows us to better appreciate lysosomal function in physiological and pathophysiological states. Furthermore, Western blot analysis and lysosomal enzymatic assays showed that the protein content and enzymatic activity of the EL fraction were as expected, with minimum contamination from other organelles. Organelle-specific quantitative proteomics approaches such as this can help progress our understanding of the role of lysosomes in atrial physiology and pathophysiology, for example by comparing protein composition from disease tissue samples or cell lines with those of healthy donors or patients.

Creation of an atrial endo-lysosomal organelle database offers valuable data in studying cardiac physiology. Using this method we identified endo-lysosomal marker proteins^31^ Rab7A, VPS29, MAN2B1, LAMTOR1, LAMTOR2, LAMTOR3, LAMTOR5, RILP, ACP2, GBA, and GAA in our proteomics data as well as LAMP2 from Western Blot data. Furthermore, statistical analysis by volcano plot of quantified protein hits revealed 564 endo-lysosomal proteins significantly enriched in the EL fraction (false discovery rate 0.05).

## Results

### Density gradient approach towards acidic organelle isolation from *C. porcellus* atria

An overview of the workflow is shown in Figure 1. In order to capture lysosomal specific proteome data in guinea-pig (*C. porcellus)* atrial frozen biopsies, we further developed a density gradient organelle isolation protocol based on previous work^5,32^. The first stage was to eliminate tissue debris and plasma membrane by brief ultra-centrifugation. The supernatant enriched in sarcoplasmic reticulum (SR), mitochondria, lysosomes, endosomes and endo-lysosomes was then separated by differential density gradient based ultra-centrifugation. The much denser organelles such as crude mitochondria with mature lysosomes and most of the SR content were separated from the soluble fraction at this stage. A high degree of purity of the endo-lysosome enriched fraction was achieved by a repeated ultra-centrifugation step. The final fraction was subjected to differential density gradient levels to partition proteins into specific compartments with respective buoyant densities.

**Figure 1.**
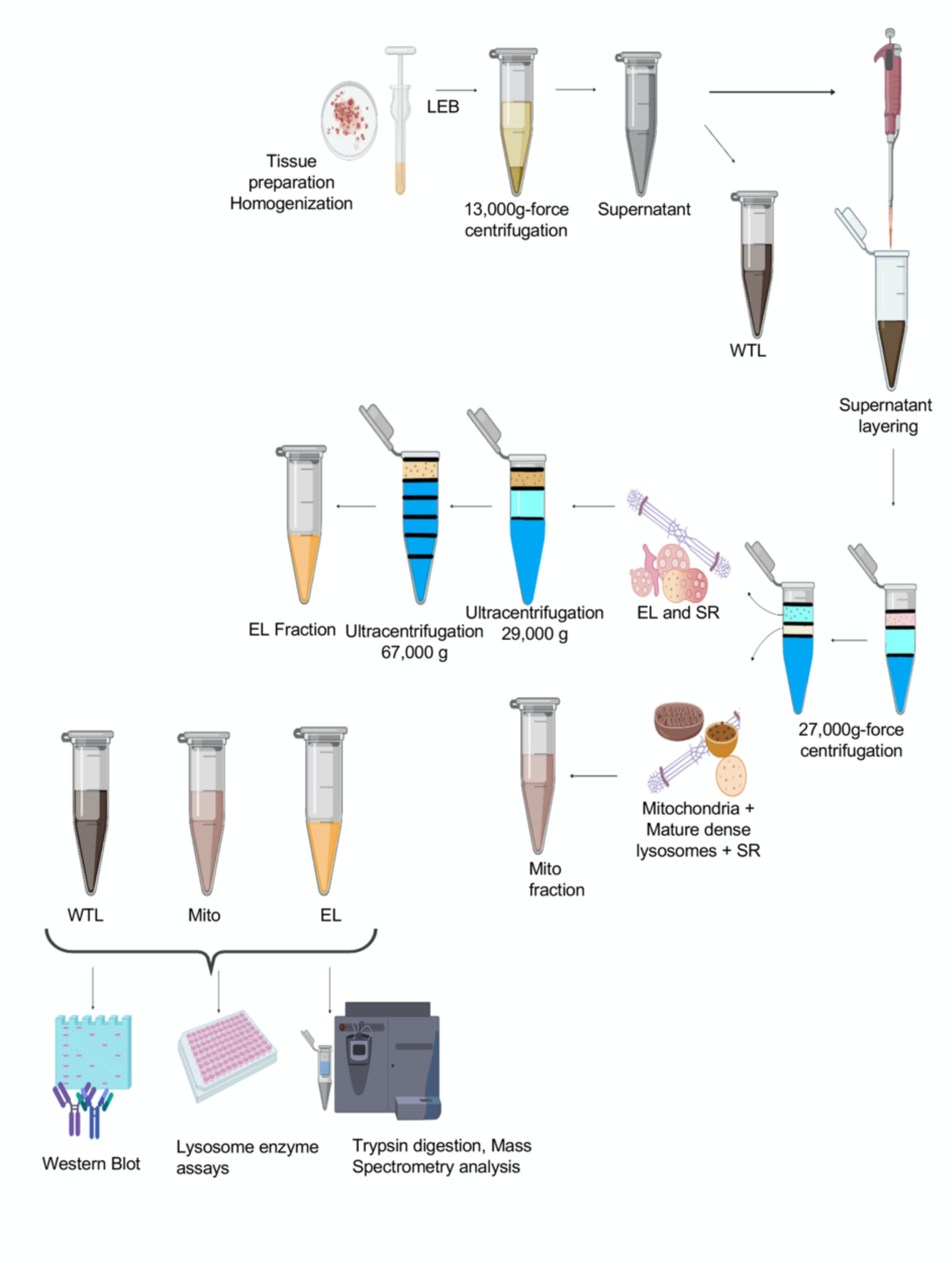
Flow chart of the acidic organelle isolation: Top: Atrial tissue is homogenized using a Dounce homogenizer; after adding Lysosome enrichment buffer (LEB), tissue lysate is briefly centrifuged at 13,000g-force, 4 °C for 2 min; supernatant is collected without disturbing the tissue pellet (tissue lysate, TL); centrifuge tube is layered with 750 μL of 2.5 M sucrose, 250 μL of Percoll, and 200 μL of TL; centrifuge at 27,000 g-force, 10 °C for 50 min: The mitochondria and SR accumulate at the bi-face of Percoll and 2.5 M sucrose and are carefully collected (mitochondria + SR enriched fraction). The area above the turbid white layer consists of endosomes and endo-lysosomes with minimum contamination of SR, and this fraction is collected for further removal of SR by centrifuging at 29,000 g-force 15 °C for 30 min; For the differential density gradient centrifugation step, centrifuge tube is carefully loaded with the different density gradients made of sucrose, Percoll and ddH_2_O (starting from 1.3 g/ml, 1.11 g/ml, 1.07 g/ml, 1.05 g/ml, 1.04 g/ml). The fraction collected from above the turbid white layer is pipetted (200 μL) on top of the 1.04 g/ml Percoll; the tube is centrifuged at 67,000 g-force at 4 °C for 30 min, and the top fraction consists of endo-lysosomes + endosomes. Bottom: Protein validation is performed using western blot, lysosome enzyme assay and proteomic analysis. (Flow chart created using Biorender.com).

### Validation of lysosomal proteins using lysosomal enzymes and immunoblotting assays

The main fractions; tissue lysate (TL), endo-lysosome (EL) and mitochondria (Mito), were validated for organelle enrichment by lysosome enzyme assays and Western Blotting (Figures 2A and 2B). Lysosomal activities of β-galactosidase and β-hexosaminidase were assayed in TL, mito, EL and the remaining gradient fractions from Dunkin Hartley guinea pig atria (Figure 2A, n = 3). Figure 2A shows total units of activity for each enzyme in each fraction. Volumes, protein amounts and specific activities for the enzymes are shown in Supplementary Data File 1.

**Figure 2.**
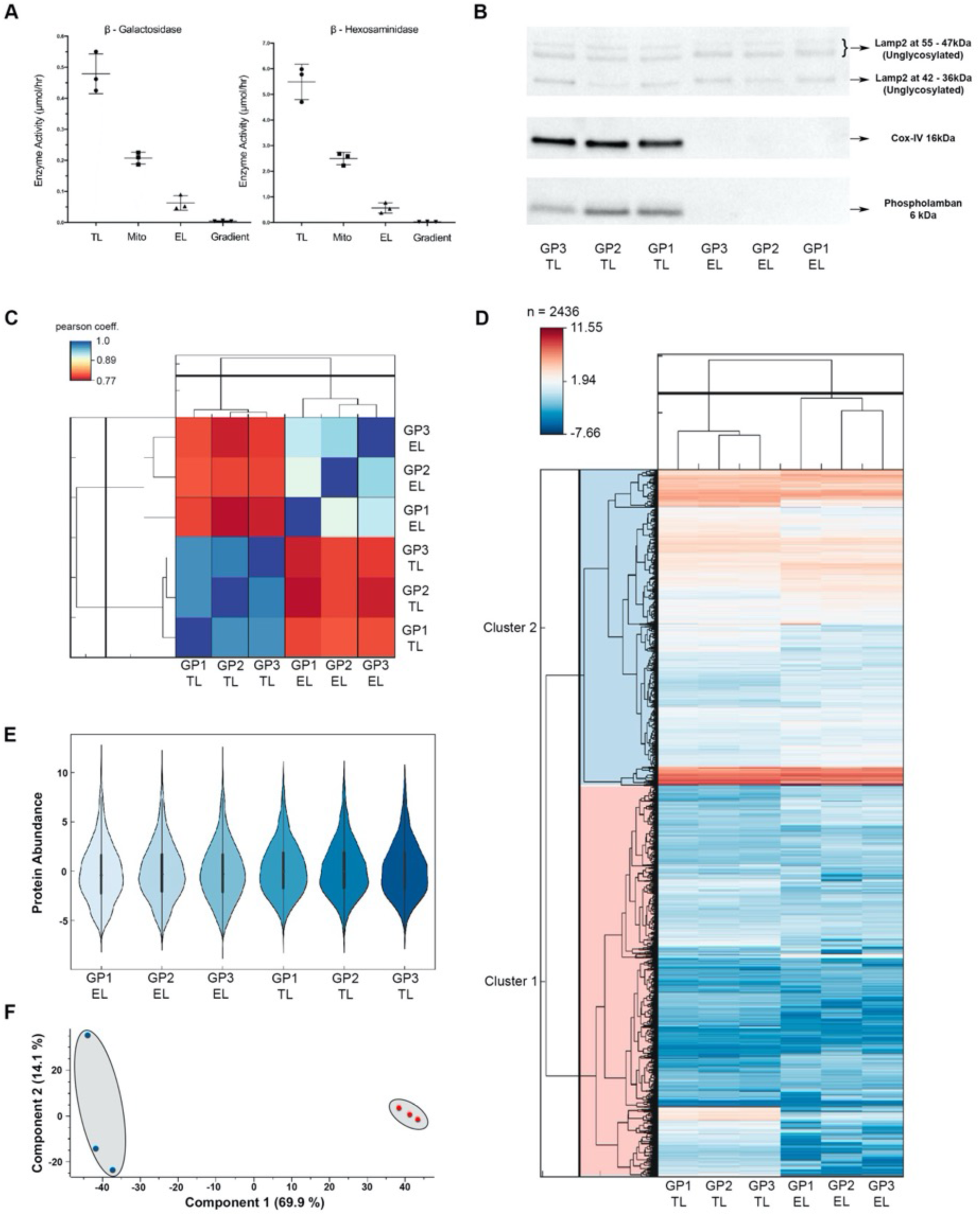
Expression of protein abundancy & technical reproducibility: **A**, Beta – galactosidase and Beta – hexosaminidase enzyme activities in adult guinea pig atria (n = 3). Lysosomal hydrolase activities (total Units) were measured in EL, Mito (which contain dense lysosomes) and as well as TL using artificial 4-MU-substrates. **B**, Western blots performed in guinea-pig atrial tissue: Lamp2, CoxIV and Phospholamban (n=3). Identification and quantification of lysosome, mitochondria and SR organelle levels between (TL) and EL fraction. **C**, Pearson co-efficient correlation plot values show the positive or direct correlation between the reliability of the triplicated samples. **D**, Heat map of z-scored protein abundances (LFQ intensities) of the differentially expressed proteins after unsupervised hierarchical clustering. **E**, Violin plot shows distribution of peptide abundance from EL fraction to TL among the triplicates. **F**, Principal component analysis (PCA) of the six atrial samples based on their proteomic expression profiles. Each data point represents the total protein groups in each sample as a single vector. The components 1 and 2 represent the spatial resolution among the vectors. The average of vectors corresponds to a point in the K-space. Component one explains 69.9% of the variation, component two 14.4%. Red: TL, Blue: EL. Panels **C**-**D** generated using Perseus 1.5.2.4 and redrawn using Instant Clue^109^.

Western Blotting was conducted to analyse the presence of EL membrane proteins by blotting against LAMP2 (Figure 2B). The absence of the predominant organelles such as SR and mitochondrial membranes were examined by the SR marker protein Phospholamban and inner-mitochondrial membrane marker protein COX IV. We observed clear band visibility of LAMP2 in all the biological replicates of EL and TL. Phospholamban and COX IV were negligible in EL (Figure 2B and Supplementary Figure 2). In addition, western blotting was carried out using Glyceraldehyde 3-phosphate dehydrogenase (GAPDH) as a loading control (Supplementary Figure 2B).

Sample intensity distributions showed high similarity, with Pearson correlation coefficients of >0.9 (TL) and >0.8 (EL) for intra-group comparisons (Figure 2C), with density histograms of the LFQ intensity data assuring the near normal distribution of protein intensities between the TL and EL in three biological replicates showing technical reproducibility (Supplementary Figure 1).

### Differential protein distribution using quantitative proteomic analysis

For overall assessment of functional protein resemblance between the fractions, unsupervised hierarchal clustering was employed on 2436 *C. porcellus* proteins. Gene ontology annotations identified statistically different abundance of protein groups between the fractions (FDR<0.05). A heat map was generated (Figure 2D), where colour representation from blue to red demonstrates lowest to highest relative abundance of the protein groups in EL compared to TL. The protein groups represented in white showed no significant difference between the protein intensity leading to different levels of expression in the protein groups. The differential distributions of protein abundance from EL to TL are demonstrated using violin plots in Figure 2E. Principal component analysis (PCA) reduced the data dimensions for simpler interpretation (Figure 2F). As indicated in Figure 2F, vector deviation of 69.9% was observed between TL (blue symbols) and EL fractions (red symbols). An exceptional 14.1% segregation was observed in first biological replicate of EL. We observed higher level of mitochondrial proteins in the TL vectors compared to EL, such as proteins involved in electron transport chain (eg: A0A286XXR8/NDUFB4, H0V9U7/NDUFB9^33^) (Supplementary Data File 2) whereas EL contained proteins from endosomal and endocytic trafficking pathways (e.g.: A0A286XGA7/TOM1^34^, A0A286XNV7/ADIPOQ^35^, A0A286Y1Z7/CDC42^36^) (Supplementary Data File 3). The molecular functional differences were clearly distinguished in the PCA plot.

### Functional networks within Endo-lysosomes

Proteins demonstrating differential abundance between TL and EL fractions were identified using volcano plot (Figure 3A). Of a total of 2436 quantified proteins, 690 accounted as depleted in EL, demonstrating higher abundance in TL fractions (Figure 3A, green and Supplementary Data File 2) whereas 564 proteins accounted for the most enriched hits corresponding to higher abundance in EL (Figure 3A, red and Supplementary Data File 3). The functional networks of the quantified, statistically significant enriched proteins acquired from volcano plot analysis were mapped using several genomic and proteomic annotations (detailed description in methods section). The STRING network created by Cytoscape generated 125 proteins involved in a single functional network, and 7 proteins displayed a detached cluster that were not connected to any functional edge (Figure 3B and Supplementary Data Files 4-5).

**Figure 3.**
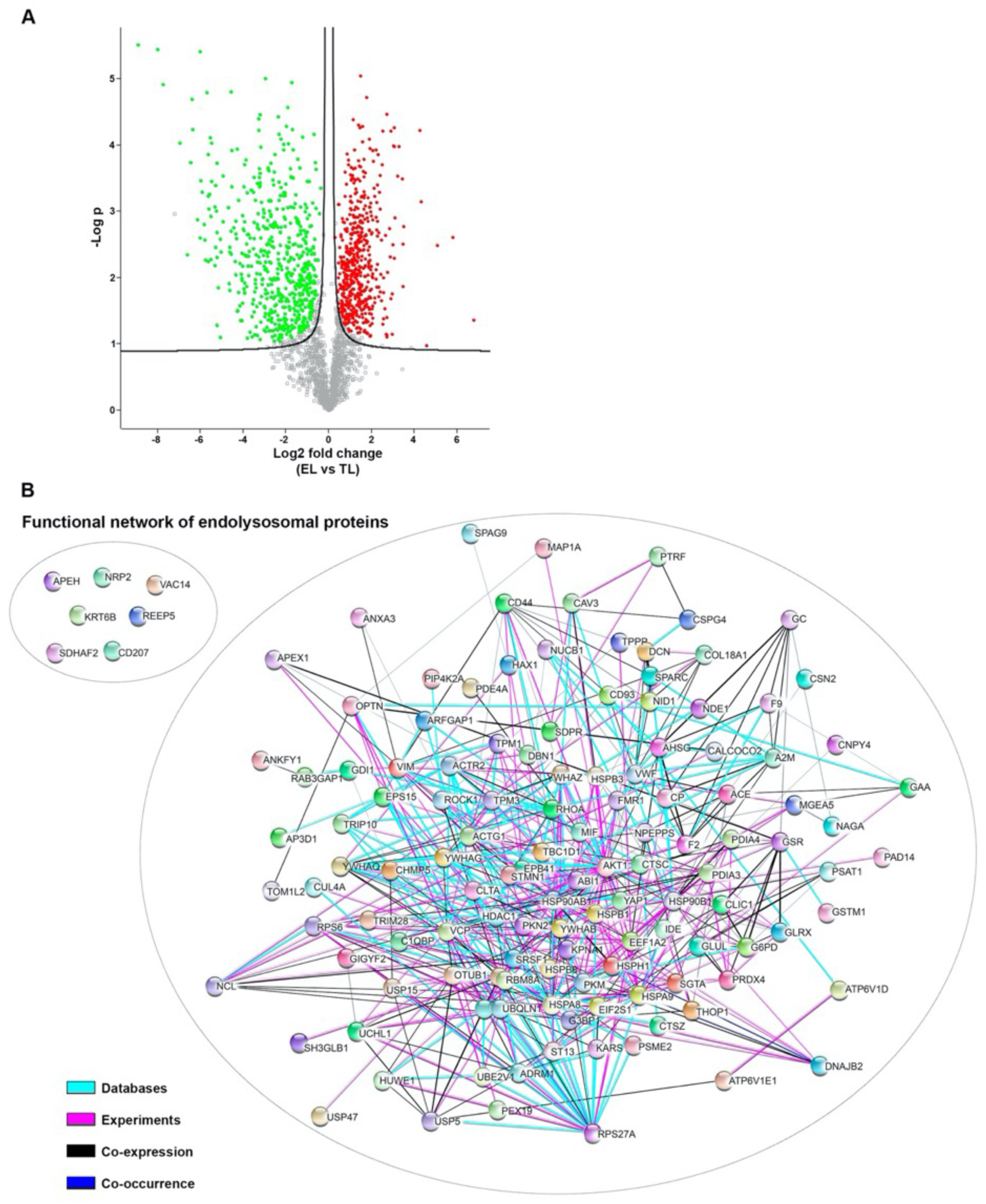
Profile of the endo-lysosomal proteins in EL fraction at a glance: **A**, Volcano plot is plotted against the −log2 transformation of the p values vs. the protein abundance differences in EL and TL. Significantly higher abundant proteins in EL compared to TL are highlighted in red and less abundant in green, respectively (FDR 0.05). **B**, EL fraction proteins were clustered using Cytoscape consortium 3.7.2 with String, KeGG, GO and Reactome pathway annotations. Proteins were clustered using median confidence score (0.4) and the molecular pathway parameters (edges) were filtered to databases, experiments, co-expression and co-occurrence.

The biological interactions of the leading network (Figure 3B) display a single network which is based on 4 functional annotations. These annotations are based on information from published databases, experiments, co-occurrence and co-expression data (www.cytoscape.org). A colour scale depicts the functional annotations, and the fading of the colour displays the strength of the evidence. In addition to the endolysosomal network analysis we performed a separate functional enrichment analysis using the gene ontology pathway analysis; the most abundant protein IDs of the EL fraction were converted to human protein IDs and uploaded to the GO analysis (ShinyGO v0.61) within the cell component filteration. The 10 most significant enrichment of the cell component proteins were identified. The highest number of protein IDs’ (188) belonged to vesicle trafficking suggesting the identified EL proteins are localized to the endolysosome route of the cell^37,38^ (Supplementary Data File 6).

### PANTHER gene annotation pathway analysis

The protein profile of EL (Figure 4A) and TL (Figure 4B) were analysed using the PANTHER pathway database (www.pantherdb.org) using a total of 1254 quantified proteins (Supplementary Data File 6). The most enriched EL fraction proteins (564) and the proteins most abundant in TL (690) were plotted with mapped gene identifiers, using pie charts to demonstrate the overall representation of protein groups by molecular function categories. The EL fraction and TL displayed respectively; 39% and 42.8% of the catalytic activity. The catalytic representation of the EL and TL were categorized according to the hydrolase enzyme groups. The percentages between EL and TL were respectively as follows; peptidase (29.9%, 19.4%), hydrolase acting on carbon-nitrogen (but not peptide) (4.8%, 8.6%) and hydrolase acting on ester bonds (23.8%, 14%), hydrolase acting on acid anhydrides (25%, 51.6%), hydrolase activity acting on glycosyl bonds (2.4%, 2.2%), hydrolase activity, acting on acid phosphorus – nitrogen bonds (4.8%, 4.3%). Lysosome specific cathepsins were highly abundant in the peptidase category found in the EL fraction. In contrast, cytochrome oxidases of mitochondrial origin were highly abundant in TL.

**Figure 4.**
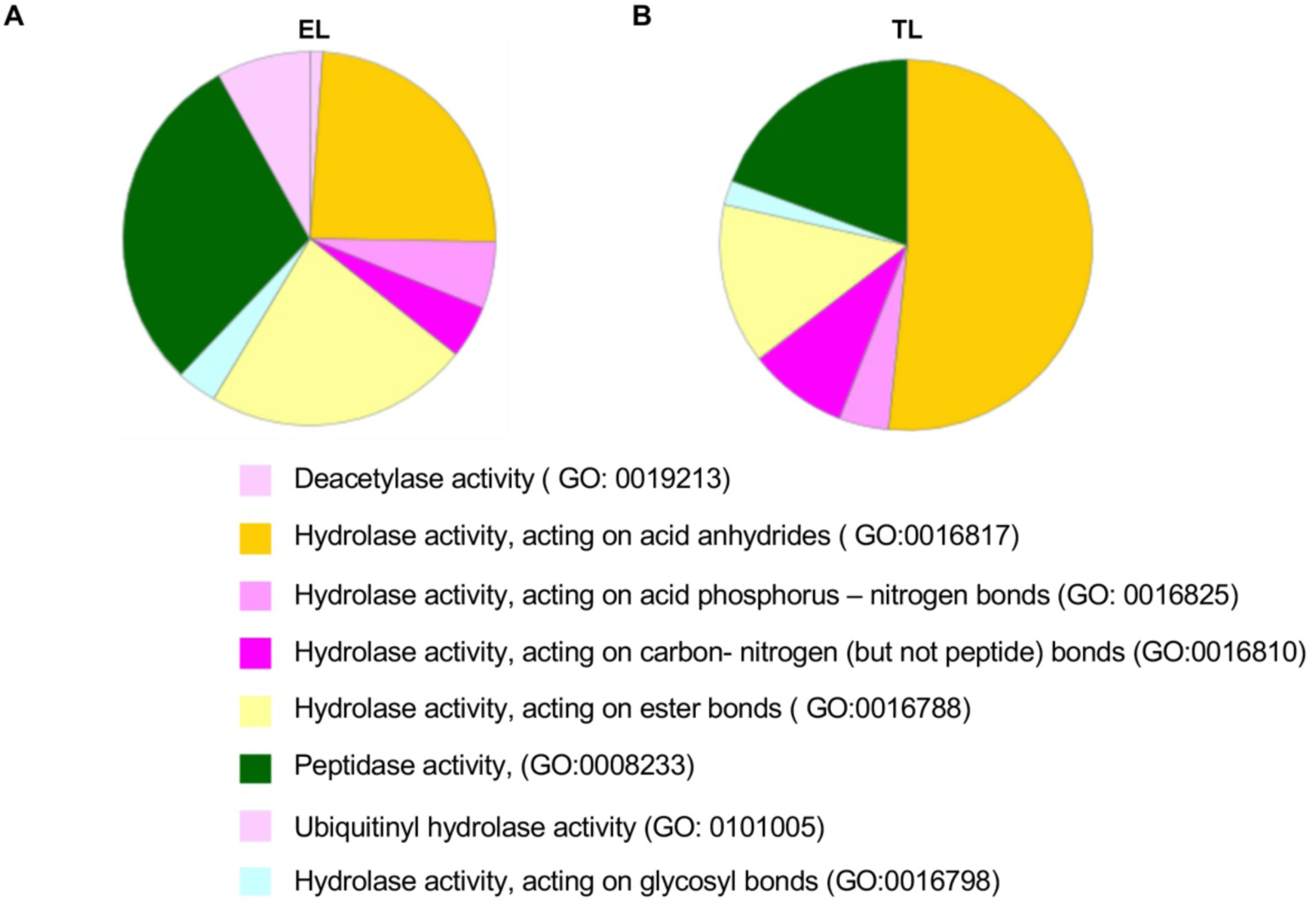
Lysosomal enzymes and immunoblotting assays: **A** & **B**: Gene ontology panther pathway analysis: The molecular function of the endo-lysosome fraction showed 39% of catalytic activity, whilst the molecular function of the tissue lysate (TL) showed 42.8% of catalytic activity. The catalytic hydrolase activity was further analysed for individual hydrolase activity, EL fraction (**A**) showed higher peptidase activity compared to TL (**B**).

### Organelle expression profile

Our data demonstrate the presence of lysosomal markers β-galactosidase, β-hexosaminidase, Rab7A, and LAMP2 in our EL preparations and the absence of markers from mitochondria (COX IV) and ER (Calnexin protein)^39^, with fewer cytochrome C reductase proteins, (ER marker; H0VNA2/CYC1, A0A286Y030/NDUFA4 in EL and A0A286XMD6/ UQCRFS1, CYC1, NDUFA4, A0A286XTF9/UQCRB, H0W408/UQCRQ, H0VIM6/UQCR10 in TL)^40^. Key proteins identified are summarised in Supplementary Table 1.

### Pathway analysis of endo-lysosomal, endosome and lysosomal proteins

The majority of the proteins identified had a direct connection with the structural biology of endo-lysosomes, lysosomes and functions related to their membranes, whilst some proteins identified had a functional role in lysosome biogenesis and endocytic processes. The following sections provide details of some of the key proteins identified in our study as well as their known associations according to previously published data.

#### a) Proteins involved in endosomal/lysosomal functions

We identified multiple proteins linked to endocytosis, including AP2A1, LRP1 and APOE^41^. AP2A1 is a key regulator of endosomal/lysosomal protein sorting pathways^42^, whereas LRP1 and APOE are involved in cholesterol metabolism^43^. H0UWL7/ACE and H0VDM6/ITGB1 have been recognized in hypertrophic cardiomyopathy disease pathways^44,45^. HSPA8 is linked to Parkinson’s disease where it is involved in the impairment of lysosomal autophagy. HSPA8 may contribute to lysosomal storage disorders (LSDs) as a functional component of lysosome vesicle biogenesis^46^. A large number of Rab proteins were categorized under Rab-regulated trafficking, membrane trafficking and endocytosis (Supplementary Table 1). We found numerous proteins, such as V-type ATPase proton pumps commonly expressed in lysosomes and lysosome related organelles, including ATP6V1D (A0A024R683)^37^ (Supplementary Table 1). In addition, we identified the endosomal proteins Flotillin-1 (A0A286XE27) ^37^ and CHMP5 (Q9NZZ3)^38^ in our EL fractions.

Atrial tissue-specific protein markers such as myosin heavy chain 6 (MYH6/ A0A286Y2B6), myosin heavy chain 7 (MYH7B/H0V2M2), peptidylglycine alpha amidating monooxygenase (PAM/H0VJZ4) and natriuretic peptide A (NPPA/H0VXX0), reported in previous cardiac proteomic studies^11^ were identified in our TL (Supplementary Table 1).

#### b) Acidic organelle proteins related to cellular Ca^2+^ and ion channels

EEA1, a phosphoinositide binding domain, and β1-integrin which has a functional role related to two-pore channels (TPC), were observed in our proteomic profiles^47,48^. Inhibition of TPC function in metastatic cancer cells has been shown to prevent trafficking of β1-integrin, leading to accumulation within EEA1-positive early endosomes and preventing cancer cell migration^47^.

EH domain-containing proteins function as retrograde transport regulators and retrograde trafficking mediates the transport of endocytic membranes from endosomes to the trans-Golgi network^49^. Our study identified several proteins involved in retrograde transport and trafficking including EHD 2 and 4, Ankyrin, Vps 35 and Annexin A. A 2010 study by Gudmundsson *et al*.^50^ showed that EHD1-4 directly associate with Ankyrin, providing information on the expression and localization of these molecules in primary cardiomyocytes and demonstrating that EHD1-4 are coexpressed with ankyrin-B in the myocyte perinuclear region. Significant modulation of EHD expression follows myocardial infarction, suggesting that EHDs may play a key role in regulating membrane excitability in normal and diseased hearts^50^. Retrograde transport is important for many cellular functions, including lysosome biogenesis where Vps35, a subunit of retromer, interacts with the cytosolic domain of the cation-independent mannose 6-phosphate receptor to mediate sorting of lysosomal hydrolase precursors to endosomes^51^. Annexin A2 another protein involved in acidic organelle Ca^2+^ binding and the endocytic pathway, is capable of active Ca^2+^-dependent plasma membrane resealing in vascular endothelial cells^52^, interacts with the lysosomal *N*-ethylmaleimide-sensitive factor attachment protein receptor (SNARE) VAMP8 and facilitates binding of VAMP8 to the autophagosomal SNARE syntaxin 17 to modulate the fusion of auto-phagosomes with lysosomes^53^.

Dysferlin (DYSF), acts as a Ca^2+^ sensor in Ca^2+^-triggered synaptic vesicle-plasma membrane fusion in myocytes^54^. Mutations in dysferlin cause limb-girdle muscular dystrophy type 2B (LGMD2B) due to defective Ca^2+^-dependent, vesicle-mediated membrane repair^55^. Loss of dysferlin causes death of cardiomyocytes, notably in ageing hearts, leading to dilated cardiomyopathy and heart failure in LGM2B patients^56^. These observations in conjunction with our data demonstrate the need for further studies related to the role of dysferlin in cardiac atrial pathology.

Annexin 6, involved in Ca^2+^ binding to cellular membranes such as those of acidic organelles including late endosomes^57^ were identified from our proteomic studies. As a regulator of the apical membrane events of the placenta, Annexin 6 binds in both a Ca^2+^-dependent and in a Ca^2+^-independent fashion^58^. Annexin 6 is also involved in the trafficking events between endocytic compartments and lysosomes leading to degradation of low density lipoproteins (LDL)^59^.

### Cross-species comparison of *C. porcellus* atrial and lysosomal proteome with the human proteome

Following conversion of the protein identifiers of the *C. porcellus* atrial proteome, protein identities were compared to previously published data for the human proteome^11,60^ for both TL (Supplementary Figure 3) and EL (Supplementary Figure 4) fractions. As shown in Supplementary Figure 3, 22.2% of the proteins identified in our *C. porcellus* TL fraction have been previously identified as showing differential expression with upregulation in the human atrial proteome when compared to ventricles^11^. These genes were spread evenly throughout the *C. porcellus* atrial proteome when the range of identified proteins was sorted according to expression level (Supplementary Figure 3A). Supplementary Figure 3B shows the most highly expressed 10% of proteins identified in the *C. porcellus* atria from our data. 20.6% of these proteins were previously identified as showing upregulation in the human atrial proteome^11^. Several genes from this group, including the most highly expressed protein from our data, myosin heavy chain α (MYH6), as well as clathrin heavy chain 1 (CLTC), talin-1 (TLN1) and heat shock protein HSP90B1, have previously been shown to demonstrate increased expression within the human atrial, compared to the ventricular, proteome^11^.

In contrast to the data in Supplementary Figure 3, which shows comparison of our TL data with proteins differentially expressed in human atria compared to ventricles, the comparison in Supplementary Figure 4 compares proteins identified in our *C. porcellus* EL fractions with the complete proteomic data for human lysosome reported by Tharkeshwar *et al.*^60^. 50.5% of genes identified in the *C. porcellus* EL proteome matched those previously identified in the human lysosomal proteome^60^ (Supplementary Figure 4A). This expression overlap increased to 66.4% of the 10% most highly expressed genes in the *C. porcellus* EL proteome (compared with all genes identified in human lysosomes^60^). Genes within this top 10% that were identified in both species included aconitase type II (ACO2), actin alpha cardiac muscle 1 (ACTC1) and the heat shock proteins HSPD1, HSP90AA1 and HSPA9 (Supplementary Figure 4B). Comparisons of proteomic data previously published from human atrial tissue^11^ and data from *C. porcellus* from our study, including relative overlaps in expression between TL and EL fractions, are shown in Supplementary Figures 5 and 6.

## Discussion

In this manuscript, we present a modified density gradient method of endo-lysosomal organelle isolation suitable for use with frozen tissue samples of at least 100mg. After confirming the identity and purity of these fractions using enzyme activity assays (Figure 2A) and Western Blots (Figure 2B and Supplementary Figure 2), we performed label-free LC-MS/MS peptide analysis and present what we believe to be the first comprehensive dataset of *C. porcellus* endo-lysosomal focussed organelle proteomics.

The importance of lysosomes for cardiac protein turnover and the role that changes in these organelles have to this function has been known for several decades^18^. In spite of this, lysosomes have remained relatively understudied in terms of a possible role in cardiac pathogenesis. Several lysosomal storage diseases may present with cardiac abnormalities as part of disease progression, e.g. hypertrophy and conduction dysfunction in Anderson-Farbry’s disease^61,62^. More recently, a role for lysosomes and endo-lysosomes as acute signalling organelles has been identified in a number of cell types^63^ including both atrial^16,64^ and ventricular^15,16,65^ cardiomyocytes. These observations spark a renewed interest in the function and protein composition of lysosomes and endolysosomes in this tissue. However proteomic databases of cardiac lysosomes have not been published to this point.

The involvement of lysosomes in cardiovascular disease has been of interest for decades. Early observations presented in patients undergoing open-heart surgery suggested that the number of lysosomes in the right atrium is increased in patients with atrial septal defects^23,24,66^. Following these observations, in 1967, Kottmeier and Wheat pursued experiments to see if similar findings could be produced in an experimental model^24^. They found that the number of lysosomes increased significantly following the creation of atrial septal defects in dogs, with the most marked increase occurring in the right ventricle providing early evidence to support the role of the lysosome as an important intracellular organelle which is related to cellular stress.

Cardiomyocytes are responsible for the beat of the heart and make up the bulk of cardiac tissue by volume^67^. These cells are structurally specialised for excitation-contraction coupling, containing large numbers of contractile filaments and mitochondria by volume. Although cardiomyocytes dominate heart tissue volume, non-myocytes (eg. fibroblasts, endothelial cells, vascular smooth muscle) are greater by nuclear number^68,69^. Consequently, whole tissue analysis of the cardiac proteome is dominated by contractile, mitochondrial and cell/ECM structural proteins. For instance, Doll *et al.* (2017) found 25% of identified protein molecules were from just six proteins in human heart tissue samples, of which two were contractile and two structural^11^. Robust conclusions regarding the effects of physiology and disease on the proteome of other organelles therefore requires accurate enrichment of the organelle of interest. An elegant method to purify endo-lysosomes from cultured HeLa cells utilising superparamagnetic iron oxide nanoparticles (SPIONs) was published in 2017^60^. A similar approach utilising the uptake of latex beads was shown to allow endosome purification in cultured macrophages^32^. Both of these methods, however, rely upon the uptake of particles to live cells, limiting their utility for analysis of frozen samples, such as might be available from large-animal and/or patient biopsies for clinical cardiac projects. The LOPIT method^9^ on the other hand, requires proteomic runs of a large number of cellular fractions, making it prohibitively expensive for smaller research groups. Instead, we focussed on improving the specificity and purity of samples produced by density-based fractionation.

Lysosomes and endo-lysosomes at different stages of their maturation pathway show a wide variety of densities which overlap markedly with other cellular organelles^5,7,70,71^. By allowing the densest lysosomes to be collected with the mito fraction, we have been able to separate a highly purified EL fraction at a density of 1.04 g/ml containing over 1200 identifiable proteins by LC-MS/MS analysis (see Supplementary data and PRIDE database entry PXD021277). Isolation of this fraction from three separate frozen *C. porcellus* atrial tissue preparations clearly established the technical reproducibility of our method, as indicated by principal component and correlation analyses (Figures 2C and F). Fig 2A shows the average (± SD) total enzymatic activity of both for β-galactosidase and β-hexosaminidase when compared to either TL or mitochondrial fractions (Figure 2A in each fraction). Although there was only about 10% of each enzyme activity in the EL fraction, contamination from either mitochondria or SR, (which can also be found in a range of densities after tissue homogenisation)^5^ was minimal and confirmed by Western Blot analysis, showing that the EL fraction was devoid of COX IV and phospholamban staining respectively (Figure 2B and Supplementary Figure 2). The mitochondrial fraction, which contains mature dense lysosomes and the denser endo-lysosomes, retained 56% of the total enzyme activity. Assay of the pooled remaining gradient fractions contained less than 0.5% of the total activity. That we recovered only 67% of the total activity of each enzyme was likely to be because of the very low protein concentration in the EL fractions (∼100µg/ml) (see the table Supplementary Data File 3). Quantitative proteomic comparisons of TL and EL from three cardiac atrial *C. porcellus* fractionations demonstrated enrichment of endosomal and endocytic trafficking pathways at the expense of mitochondrial proteins such as components of the electron transport chain. In particular, we identified enrichment of a range of known lysosomal markers within the EL fraction: β-galactosidase, β-hexosaminidase, β-glucosidase, Rab7, and LAMP2.

Comparison of the proteins identified using our fractionation method to previously published results^11,60^ provides validation of the method presented in this manuscript. Just over half of genes associated with the proteins found in our study were also seen in human lysosomal/endo-lysosomal samples from SPION-aided isolation^60^ (Supplementary Figure 4). This rose to nearly two-thirds of the most highly expressed proteins (Supplementary Figure 4B). The percentage of matching proteins is lower in our TL samples when comparing to human atrial samples^11^, however, still includes the most highly abundant protein in our dataset, myosin heavy chain alpha (MYH6), as well as other abundant proteins. Importantly, our dataset also includes proteins known to be abundant within atrial tissue (eg. RYR2). RYR2 is the major calcium release channel found on the cardiac SR and mediates calcium-induced calcium release (CICR). The inclusion of this, and other cardiac membrane proteins within our dataset (e.g. Sarcoplasmic/endoplasmic reticulum calcium ATPase 2 (SERCA2), Sodium/calcium exchanger 1, Sarcolemmal membrane-associated protein, Sarcoglycans, Sarcoplasmic reticulum histidine-rich calcium-binding protein, G protein-activated inward rectifier potassium channel 4, Anion exchange protein 3, Voltage-dependent calcium channel subunit alpha-2/delta-1, Dystrophin and Plasma membrane calcium-transporting ATPase 4 (PMCA4)), is encouraging as identification of membrane-spanning proteins requires robust cell lysis and homogenisation.

Our EL proteomic data identified the presence of multiple protein hits relevant to diseases that have been linked to dysfunctional lysosomal enzymes. These include lysosomal α-glucosidase (GAA), a key lysosomal enzyme involved in the degradation of glycogen in lysosomes^72^, the lysosomal protective protein/cathepsin A/H0VMB1, which serves a protective function by regulating stability and activity of beta-galactosidase and neuraminidase enzymes^73^ and also plays a role in galactosialidosis^74^, Clusterin/H0VVP2, identified as a potential biomarker for the lysosomal storage disorder mucopolysaccharidosis^75^, and Decorin, a protein that when dysregulated contributes to cardiac fibrosis or fibrotic stiffness^76^. Glycogen phosphorylase, brain form (PYGB), is a lysosomal enzyme identified in our EL fraction that regulates glycogen mobilization^77^, and plays a prominent role as the only marker protein elevated in the early-stage of asymptomatic patients with Fabry disease^78^. In addition, we identified a major complement of lysosomal cathepsins in our proteomic data that have been previously linked with cardiovascular diseases, including cathepsins B, C, D and Z^79–85^. The successful identification of such disease related proteins highlights the potential for use of these techniques in understanding the role of lysosomal proteins in pathology.

Our EL preparations demonstrated the enrichment of lysosomal markers β-galactosidase, β-hexosaminidase, Rab7A, and LAMP2. Apart from general lysosomal proteins, EL fractions highlighted cardiac specific lysosomal proteins such as putative phospholipase B-like 2 (PLBD2)^86^ whereas TL fractions highlighted cardiac specific muscle troponin (TNNT2). Increased plasma troponin level is a risk stratification factor in AF for myocyte injury and also in myocardial infarction ^87,88^. The atrial dilatation factor natriuretic peptide (as N-terminal pro B-type natriuretic peptide), can be determined in serum/plasma samples and levels compared depending on AF presence and mode of detection^89^. We identified ANP in both of our TL and EL AF^90^.

Detection of proteins specific to the atria, such as Natriuretic peptides A (NPPA/ ANP), highlight the utility of such methods to study atrial cardiovascular diseases. The *NPPA gene* is expressed primarily in the heart, where the expression level is higher in atria than ventricles^91^.

An important protein we detect in the EL fraction is Rab7, a small GTPase that belongs to the Rab family, known to control transport to late endocytic compartments such as late endosomes and lysosomes^92^. Rab7 promotes lysosomal biosynthesis and maintains lysosomal function^93^. Rab7 is directly or indirectly involved in each event that occurs between early endosomes and lysosomes. Endo-lysosomes are known to serve as intracellular iron storage organelles^94^. Fernández et al^95^ show that increasing intracellular iron causes endolysosomal alterations associated with impaired autophagic clearance, increased cytosolic oxidative stress and increased cell death and these effects are subject to NAADP. Cell death triggered by altered intralysosomal iron handling is abrogated by inhibiting RAB7A activity. Alterations in the activity of Rab7 may be associated with cardiovascular diseases, lipid storage disorders and neurodegenerative diseases^96–98^.

As mentioned in the introduction, previously studied lysosomal protein profiles such as Schröder *et al.*^99^, used liver cells for the lysosome isolations. The use of liver cells for lysosome isolation was to alter the density of lysosomes. These studies utilised injection of Triton to the animal models, which is metabolised by the liver lysosomes. Dextran accumulated hepatic lysosomes become enlarged and denser than its natural existence so that both sedimentation coefficient and equilibrium density are increased in sucrose gradient^100^. In our endo-lysosome isolation protocol, such modifications or alterations were not used to manipulate the nature of the endolysosomes.

The fractionation method presented here is able to isolate cardiac lyso/endo-lysosomes in a robust and repeatable manner. This manuscript presents, to our knowledge, the first cardiac lyso/endo-lysosomal proteomic data set. We chose *C. porcellus* for this work due to its known electrophysiological similarities to human cardiomyocytes and long-established use for physiology data collection within the field^101^. Given the interest in lysosomes and endo-lysosomes as catabolic, storage and acute signalling compartments in cardiomyocytes, the use of this method for analysis of further species and in order to compare how the pathophysiology of clinical samples and disease models affects these organelles is of great interest for future research^102,103^. Knowledge obtained from a combination of experiments performed at many levels; genes, proteins, single cells, *in vitro* tissue engineering, isolated cardiac tissue, whole organ, *in vivo* cardiac studies, rather than a single model or experimental technique, will lead to improved strategies for diagnosis and treatment.

### Limitations

The modified density gradient protocol described in this study has the potential to identify lysosomal proteins using relatively small sample volumes. However, it is important to recognise that the proteins identified using this technique are likely to be an underestimate of the total proteins present within samples. For example, our analysis did not detect TPC1 or TPC2 proteins, endo-lysosomal ion channels that would be expected to be present^104–107^. Detecting these proteins in proteomic screens is problematic primarily due to their hydrophobicity, low levels of expression and lack of trypsin cleavage sites in their transmembrane segment sites^108^. We did however observe β1-integrin, which has a functional role related to two-pore channels (TPC)^47^, in our EL fractions. Disrupted TPC function also halts trafficking of β1-integrin, leading to accumulation in EEA1-positive early endosomes^51^. More recently, the contribution of TPC2 and NAADP to acute and chronic β-adrenoceptor signalling in the heart has been demonstrated^16^. In addition, it is not possible to rule out the possibility that some small contamination of the EL fraction with early endosomal proteins may occur during the fractionation phase. For example, besides endo-lysosomal proteins, we discovered potential minor contamination from early endosomal proteins in our EL fraction such as EEA1, RAB1A, RAB6A.

## Methods

### Animals

All experiments were performed in accordance with Home Office Guidance on the Animals (Scientific Procedures) Act 1986 (UK). Hearts were swiftly isolated from six healthy Duncan Hartley male guinea pigs following cervical dislocation and immediately perfused with ice-cold heparinised phosphate buffered saline (PBS). Both left and right atria were dissected, snap frozen in liquid nitrogen and stored at −80° C until required.

### Tissue homogenization

Frozen atrial tissue biopsies (100mg) were thoroughly cleaned in PBS and weighed. A minimum of 100 mg tissue is tissue is required in order to perform proteomics and molecular biology (enzyme assay and Western Blots). Each atrium was quartered and gently homogenised using a 7 ml Dounce homogeniser in Lysosome isolation buffer (LIB) [Containing 1:500 protease inhibitor cocktail (PIC) and phosphatase inhibitor (PHI) (Bio vision), (PhosSTOP Roche)]. Preparations were further homogenised in 1 ml Dounce homogeniser and transferred to chilled 1.5 ml ultracentrifugation tubes (Beckmann coulter). Sample preparations were mixed at a ratio of 1:1.5 Lysosome enrichment buffer [(LEB) (Biovision, containing 1:500 PIC)] to homogenate by inverting tubes, and were stored on ice for 5 min until the centrifugation.

### Isolation of acidic organelles by fractionation

Samples were centrifuged at 13,000 g for 2 min at 4°C (TLX Beckmann Coulter Ultra Centrifuge) and the resulting supernatant, corresponding to the TL was collected. Further fractionation was processed using 75% of the collected TL. 1.5 ml ultracentrifuge tubes were underlaid with 750 µl of 2.5 M sucrose (Fisher Scientific) followed by 250 µl of Percoll (Santa Cruz Biotechnology). 200 µl TL was layered on top of the percoll layer and centrifuged at 27,000 g X 50 min at 10°C. The supernatant layer just above the turbid white, mitochondrial fraction (Mito fraction) was carefully removed, and the Mito fraction itself was collected separately. The collected supernatant was retained and repeated for a further centrifugation step at 29,000 g X 30 min at 15°C (500 µl of underlaid 2.5 M sucrose with overlaid 500 µl Percoll). The supernatant above the sucrose and Percoll intermediate was collected for further fractionation. Firstly, ultracentrifuge tubes were underlaid with 2.5 M sucrose and overlaid with a series of Percoll dilutions (1.11 g/ml – 1.04 g/ml in ddH_2_O). The ultracentrifuge tubes were centrifuged at 67,000 g X 30min at 4°C. The fraction at 1.04 g/ml was collected and labelled as the endolysosomal fraction (EL). N = 3 guinea pigs were used for western blots and proteomic analysis. A separate n = 3 biological triplicate was used for the lysozyme enzymatic analysis. The reproducibility of the fractions produced using biological triplicates can be found in Figure 2.

### Lysosomal hydrolase activity assays

Lysosomal hydrolase activities were performed in EL, Mito fractions and TL. To fluorometrically measure the lysosome enzyme levels, artificial sugar substrates containing the fluorophore 4-methylumbelliferone (4-MU) were used. For measuring β-hexosaminidase activity, 3 mM 4-MU N-acetyl-β-D-glucosaminide (Sigma Aldrich) in 200 mM sodium citrate buffer, pH 4.5 and 0.1% TritonX-100 was used as substrate. For β-galactosidase activity, 1 mM 4-MU β-D-galactopyranoside (Sigma Aldrich) in 200 mM sodium acetate buffer, pH 4.3, 100 mM NaCl, and 0.1% Triton X-100 was used as substrate. The reaction was stopped by adding chilled 0.5 M Na_2_CO_3_, and the released fluorescent 4-MU was measured in a Clariostar OPTIMA plate reader (BMG Labtech, Ortenberg, Germany) with an excitation at 360 nm and emission at 460 nm. A standard curve for free 4-MU was used to calculate the enzyme activity. Results were calculated as total Units of enzyme activity (nmol/hr) and also normalised with respect to protein content.

### Protein quantitation assay

Sample fractions EL and TL were mixed at a ratio of 1:1 with radio-immunoprecipitation (RIPA) buffer (Thermo scientific). Protein concentrations of all tissue fractions and TL were determined using the Bicinchonic acid assay (BCA Protein Assay Kit, Thermo Scientific). Bovine serum albumin was used as a protein standard, and serial dilutions were prepared from the initial stock concentration of 2mg/mL to prepare a standard curve. To achieve accuracy, protein assays were performed in triplicate. Absorbance values were measured at 562 nm. Protein concentrations were calculated by linear regression analysis.

### SDS/PAGE gel preparation and western blotting

Sample fractions EL and TL were solubilised, and proteins denatured using SDS/PAGE loading buffer (bio rad) and 2-mercaptoethanol (Sigma-Aldrich). Proteins were separated by gel electrophoresis (NW04120BOX, NuPAGE 4%-12% Bis-Tris protein gels, 20X MES buffer). The gel was transferred to nitrocellulose membrane (NC) (Bio-Rad) for protein transfer (X-cell-II blot module, Thermo Fisher Scientific). After two hours, NC membrane was incubated in 5% skimmed milk. The primary antibodies anti LAMP2 (1:500, PA1-655, Thermo fisher scientific), anti-COX IV (1:1000, Abcam, ab16056) and anti-Phospholamban (1:1000, Abcam, ab85146) were incubated. Goat anti-rabbit antibody (1:2500, Dako P0448) was used as the secondary antibody to detect the protein markers of lysosomes, mitochondria and SR, respectively. The secondary antibodies were detected via chemiluminescence using Westar Supernova (XLS3,0020, Cyanogen) and the protein bands were visualised in a ChemiDoc XRS+ imager (Bio-rad with image Lab software).

### Mass-spectrometry analysis

The samples were reduced by the addition of 5 mL of 200 mM dithiothreitol (30 minutes at room temperature) and alkylated with 20 mL of 200 mM iodoacetamide (30 minutes at room temperature) followed by methanol-chloroform precipitation. The pellets were resuspended in 6 M urea in 400 mM TrisHCl, pH 7.8. Urea was diluted to 1 M using 400 mM Tris-HCl, pH 7.8, and the proteins were digested with trypsin in a ratio of 1:50 (overnight at 37oC). After acidification to a final concentration of 1% formic acid, the samples were desalted on Sola HRP SPE cartridges (ThermoFisher Scientific) and dried down in a SpeedVac (3-6hours).

Samples were further desalted online (PepMAP C18, 300µm x5mm, 5 µm particle, Thermo) for 1 minute at a flow rate of 20 µl/min and separated on a nEASY column (PepMAP C18, 75 µm x 500mm, 2 µm particle, Thermo) over 60 minutes using a gradient of 2-35% acetonitrile in 5% DMSO/0.1% formic acid at 250nl/min. (ES803; ThermoFisher Scientific) and analyzed on a Dionex Ultimate 3000/Orbitrap Fusion Lumos platform (both ThermoFisher Scientific) using standard parameters.

Mass spectrometry data were analyzed quantitatively with the MaxQuant software platform (version 1.6.2.3), with database searches carried out against the UniProt *C. porcellus* database (UP000005447_10141). A reverse decoy database was created, and results displayed at a 1% FDR for peptide spectrum matches and protein identifications. Search parameters included: trypsin, two missed cleavages, fixed modification of cysteine carbamidomethylation and variable modifications of methionine oxidation and protein N-terminal acetylation. Label-free quantification was performed with the MaxLFQ algorithm with an LFQ minimum ratio count of 2. ‘Match between runs’ function was used with match and alignment time limits of 0.7 and 20 min, respectively. The mass spectrometry proteomics data have been deposited to the ProteomeXchange Consortium via the PRIDE partner repository with the dataset identifier PXD021277.

### Quantitative analysis and statistics

Quantitative analysis of significant differences between the protein expression of EL and TL samples and data visualisation was performed using the Perseus software platform (version 1.5.2.4) using LFQ values of biological replicates. Data for peptides where more than two values were absent from six biological replicates were excluded from the quantitative analysis and distributions of excluded values between the EL and TL fractions are represented in the Venn analysis shown in Supplementary Figure 1B. Remaining data with no more than 2 missing values were uploaded as a data matrix in Perseus with the respective LFQ intensities as main columns. The data matrix was reduced by filtering based on categorical columns to remove protein groups only identified by site, reverse decoy hits and potential contaminants. A total of 2436 proteins remained after filtering. Groups of biological replicates for EL and TL fractions were defined in categorical annotation rows. Data were log transformed (log2(x)) and normalised via median subtraction. Missing data points were imputed based on normal distribution and visualised as LFQ intensity histograms (per biological replicate) with imputed values shown separately (Supplementary Figure 1A). Principal component analysis (PCA) (Figure 2F) was performed on 100% valid values. A volcano plot was generated based on LFQ intensities applying two-way Student’s t-test to test for significant difference of protein abundance between EL and TL protein expression (Figure 3A). A permutation-based false-discovery rate (FDR) was determined with 250 randomisations and S_0_ = 0.1 (default). Out of the total 2436 proteins, 1254 proteins were accounted with 99% confidence level at 5% FDR. Of these 690 proteins were low abundant in EL compared to TL, and 564 proteins were considered highly abundant in EL with 2 to 6 log fold change (Figure 3A).

### Network analysis

The functional networks of the statistically quantified proteins acquired from volcano plot were analysed using Cytoscape (Version 3.7.2) and Panther pathway analysis software (Figure 3B). *C. porcellus* proteins were converted to *H. Sapiens* with ID mapping to identify the reviewed proteins with 100%-50% similarity in the molecular function through biological and molecular pathways. The Cytoscape 3.7.2. programme was used in order to investigate associations and relations between EL proteins and protein interaction data was extracted from the STRING, KEGG and Reactome databases. The protein match to the molecular networks aligned with medium confidence of 0.4 and STRING identified a total number of 125 proteins involved in a single functional network and 7 proteins displayed in a detached cluster within the endo-lysosomal compartment. We selected the 564 highest abundant EL fraction proteins and clustered into the main network. We applied perfused force-directed layout on the network using clusterMaker 2.8.2. Functional enrichment analysis was performed for the two clusters. Pearson correlation coefficient, heat maps, PCA and histograms (Figure 2 and Supplementary Figure 3) were produced to assess the reliable correlation between the sample triplicates. The heat maps were produced using Euclidian distance and K-mean clustering. Plots were generated using Perseus software (v. 1.5.2.4) and presented for publication using Instant Clue^109^ The protein profile plots were created using Pearson clustering. The unavailability of the characterized whole guinea pig proteome limitation was minimized by selecting 100% - 50% overlap of the proteins with the closest species in the phylogenetic tree. These selections were automatically identified by proteome databases such as Uniprot and proteomic tools Perseus, Cytoscape, string and panther pathway.

### Comparison of identified proteins with human expression data

Proteins identified in both TL and EL samples and their corresponding gene identities were compared with proteomics data for human left and right atria previously published by Doll *et al.*^11^ and lysosomes from HEK cells previously published by Tharkeshwar *et al.* respectively^60^ (Supplementary Figures 3-6). Data from all samples within each fraction were pooled and sorted according to expression level based on - 10 LogP values as determined by PEAKS 8.5 analysis (TL = 2326 genes; EL = 1219 genes). Gene identities were compared with those from published data^11,60^ to identify overlaps in expression of gene orthologues between *C. porcellus* and human samples. Dynamic range of protein expression was sorted according to the −10 LogP values for each identified protein (analysis performed using Microsoft Excel, MaxQuant and GraphPad Prism software).

## Acknowledgements

RABB is funded by a Sir Henry Dale Wellcome Trust and Royal Society Fellowship (109371/Z/15/Z) and TA acknowledges support from The Returning Carers’ Fund, Medical Sciences Division, University of Oxford. RABB is a Senior Research Fellow of at Linacre College. RAC is a post-doctoral scientist funded by the Wellcome Trust and Royal Society (109371/Z/15/Z). SJB is funded by the British Heart Foundation, Project Grant Number PG/18/4/33521. FMP is a Wellcome Trust Investigaor in Science and a Royal Society Wolfson merit award holder. DAP was funded by the Mizutani Foundation. AG is a Wellcome Trust Senior Investigator and a Principal Investigator of the British Heart Foundation Centre of Research Excellence at the University of Oxford. We would like to thank the Sitsapesan and Tammaro Groups, Department of Pharmacology, University of Oxford for access to scientific equipment.

## Author Contributions

RABB conceived and designed the study; TA and RAC developed the organelle proteomics methodology; TA, GB, RF and HK performed LC-MS/MS; TA and RAC performed western blots; TA and DP performed enzyme assays; RAC and SJB contributed intellectually to the project and drafting of the paper; SJB performed analysis and statistics on human comparative data. TA, SJB and RF created the figures. All authors contributed to writing of the paper.

## Supplemental Information

**Supplementary Table 1.**
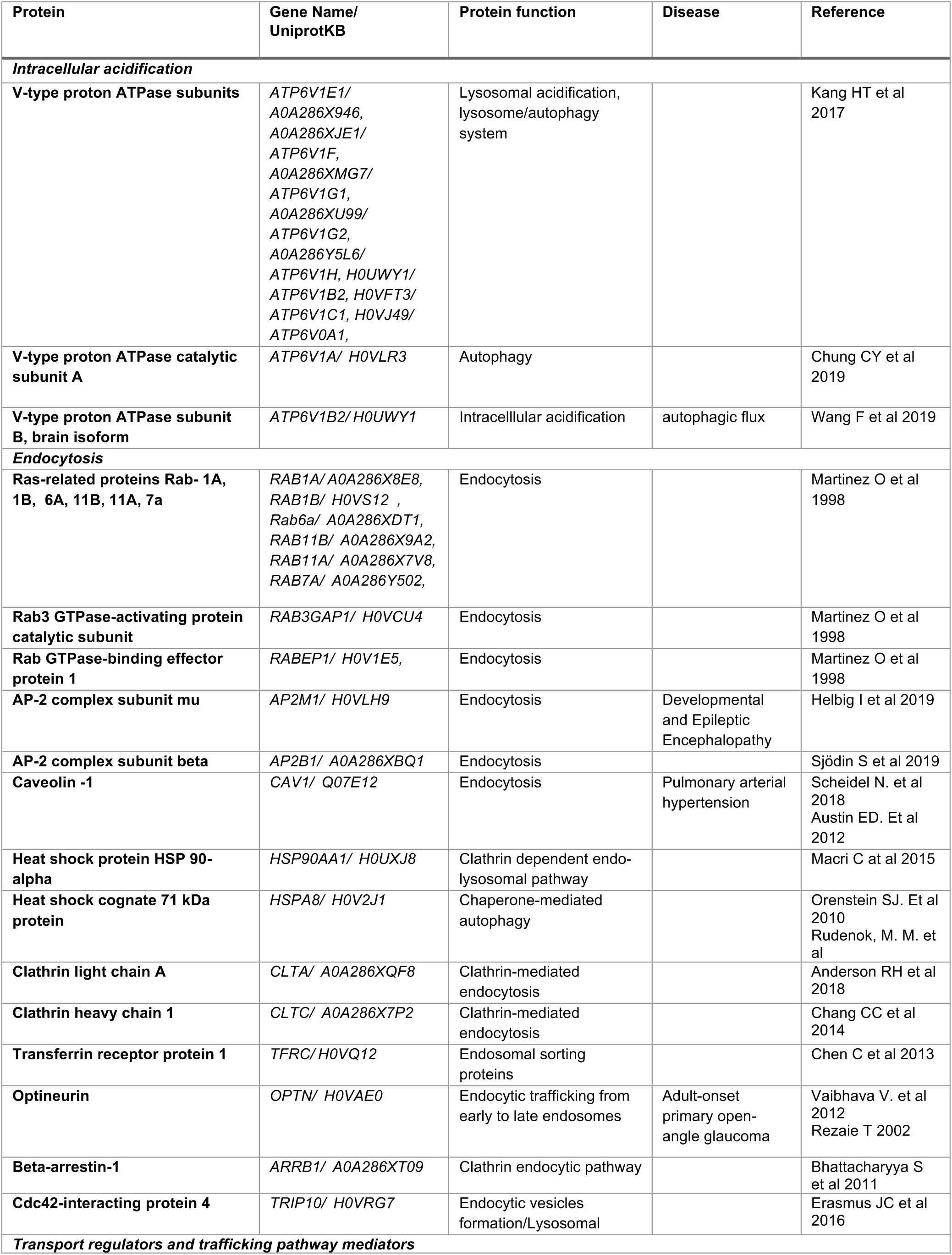
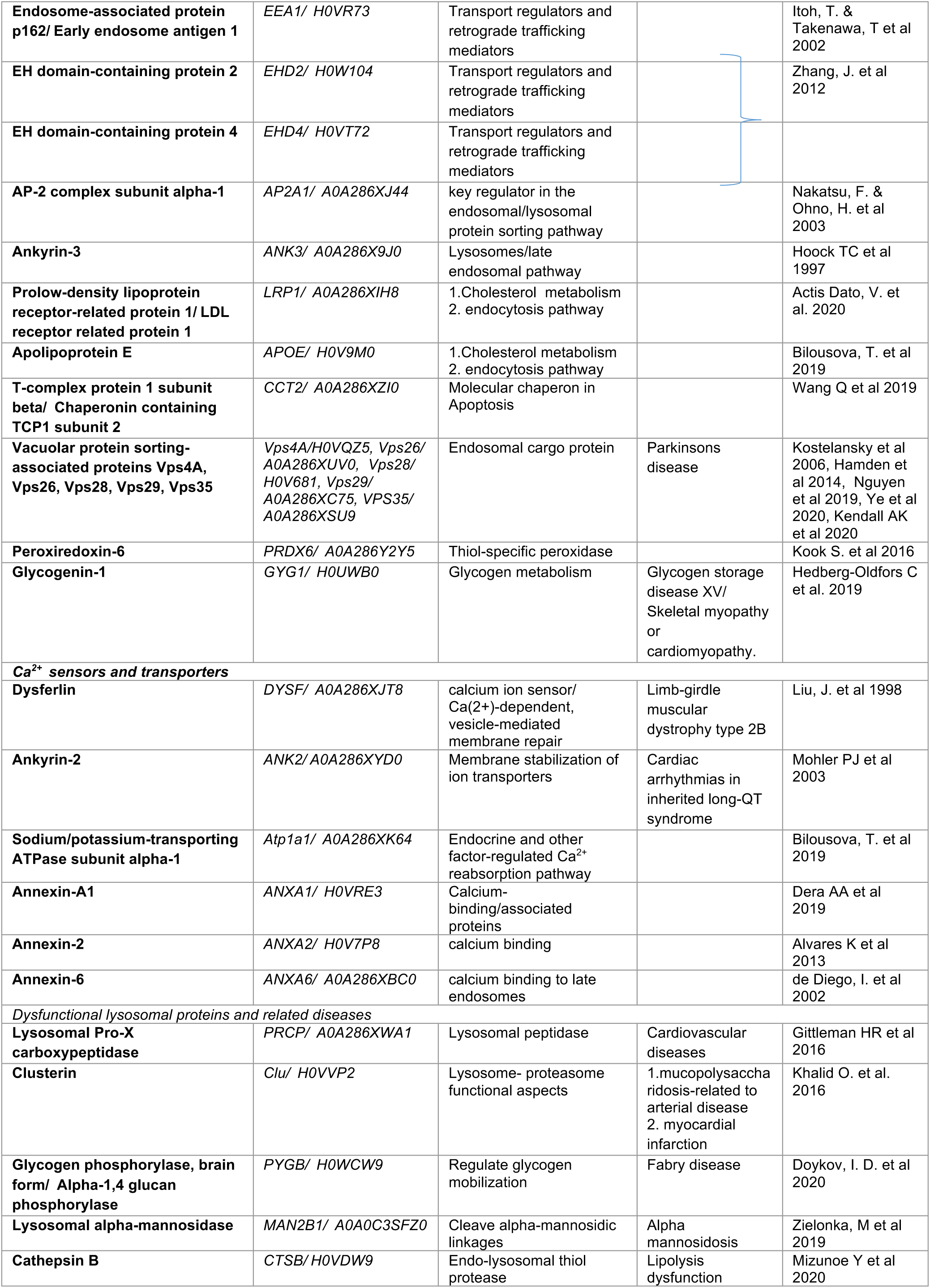
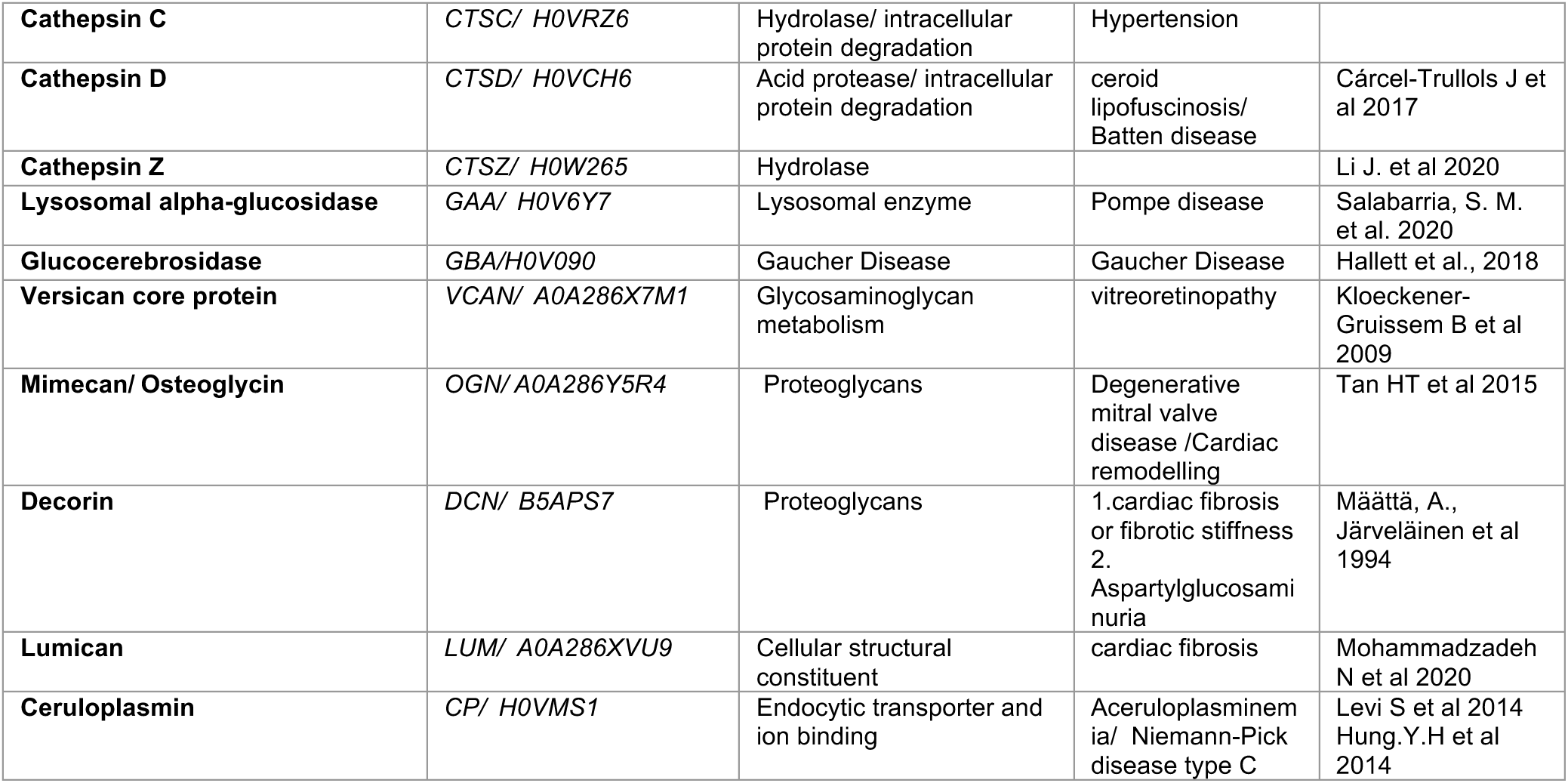
Key proteins identified from *C. porcellus* EL fraction and grouped according to protein function with information relating to their known roles in disease.

**Supplementary Figure 1.**
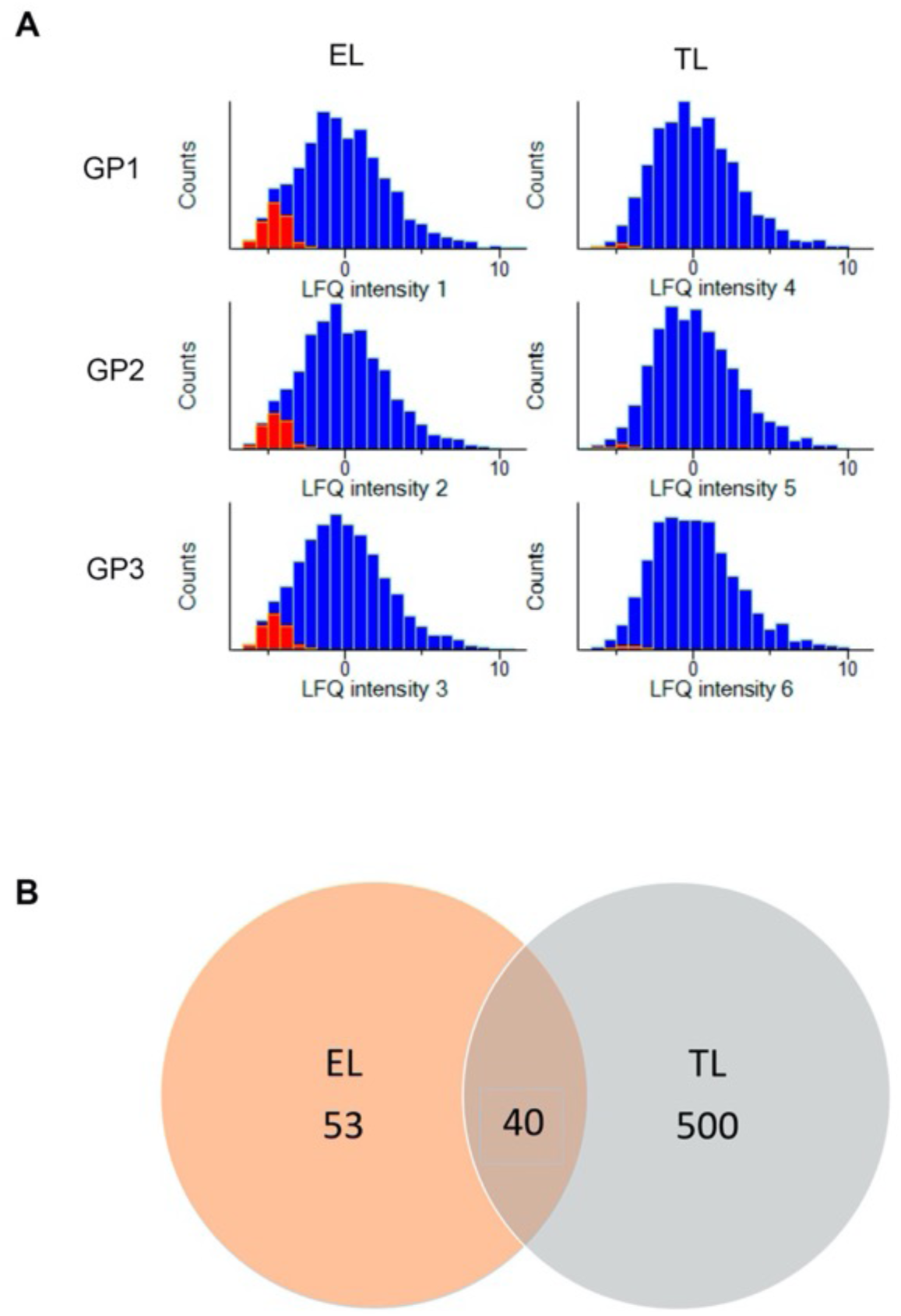
**A**, Technical Reproducibility of proteomic measurements and representation of quality control using histogram: The histogram is produced of the triplicated LFQ intensity datasets, demonstrating the normal distribution of protein intensities. Contributions of non-imputed (blue) and imputed (red) values are shown separately in histograms. Data was median subtracted for normalization. **B**, Venn diagram representing relative distribution between fractions for proteins with more than two missing values that were excluded from subsequent analysis; EL = 53, TL= 500 and 40 proteins were detected in both EL+TL.

**Supplementary Figure 2.**
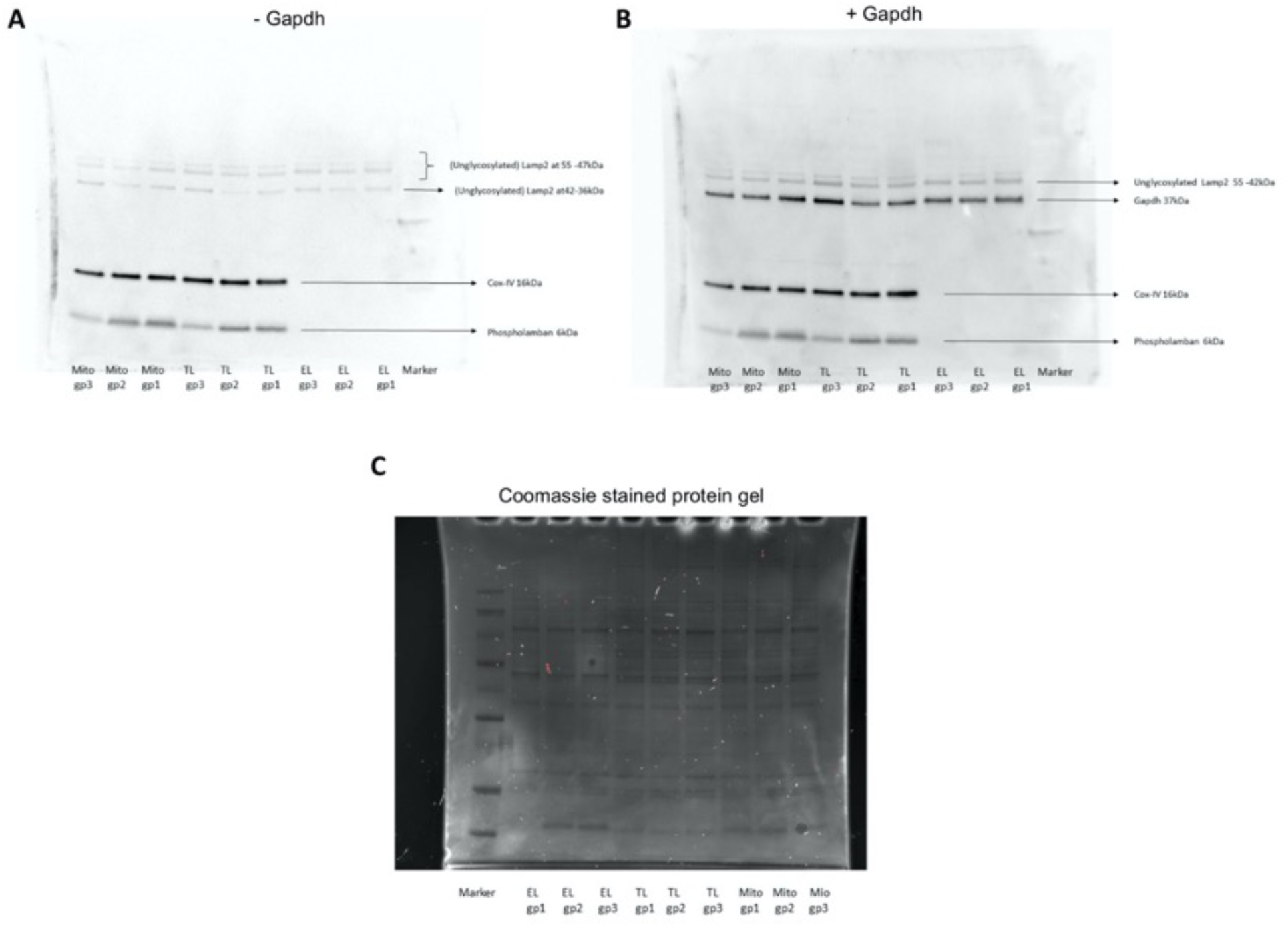
**A**, Complete Guinea pig (n = 3) western blot corresponding to Figure 2B. The western blots of the guinea pig atrial tissue used for lysosome assays displayed positive COXIV bands for the presence of mitochondria in Mito fraction and TL. EL fractions did not display bands for COXIV. **B**, Guinea pig (n = 3) western blots including Gapdh loading control. **C**, Coomassie stained gel demonstrating protein loading.

**Supplementary Figure 3.**
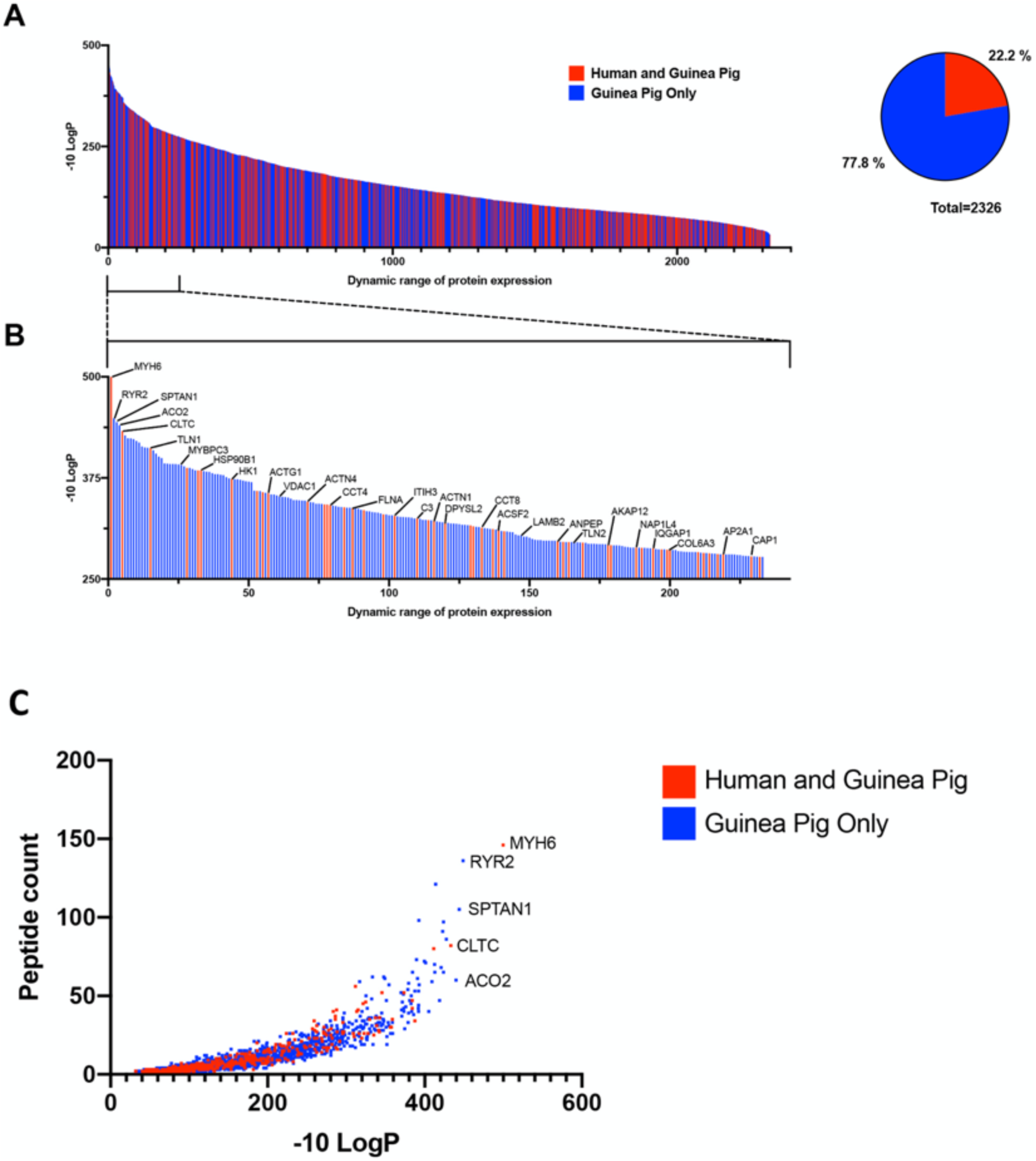
**A**, Dynamic range of peptides identified in guinea pig tissue lysate arranged according to −10 LogP values for peptides. Red bars indicate matches with genes confirmed in human heart tissue (data reproduced with permission from Doll et al.^11^). Total genes identified = 2426. **B**, expanded view of the top 10% of genes shown in A. Individual gene identifiers have been labelled for reference. **C**, Scatter plot to show relationship between −10 LogP values for all genes identified in guinea pig tissue lysate and the total peptide count for each gene. Red boxes indicate matches with genes confirmed in human heart tissue (data reproduced with permission from Doll et al.^11^). Labels indicate the most highly expressed genes.

**Supplementary Figure 4.**
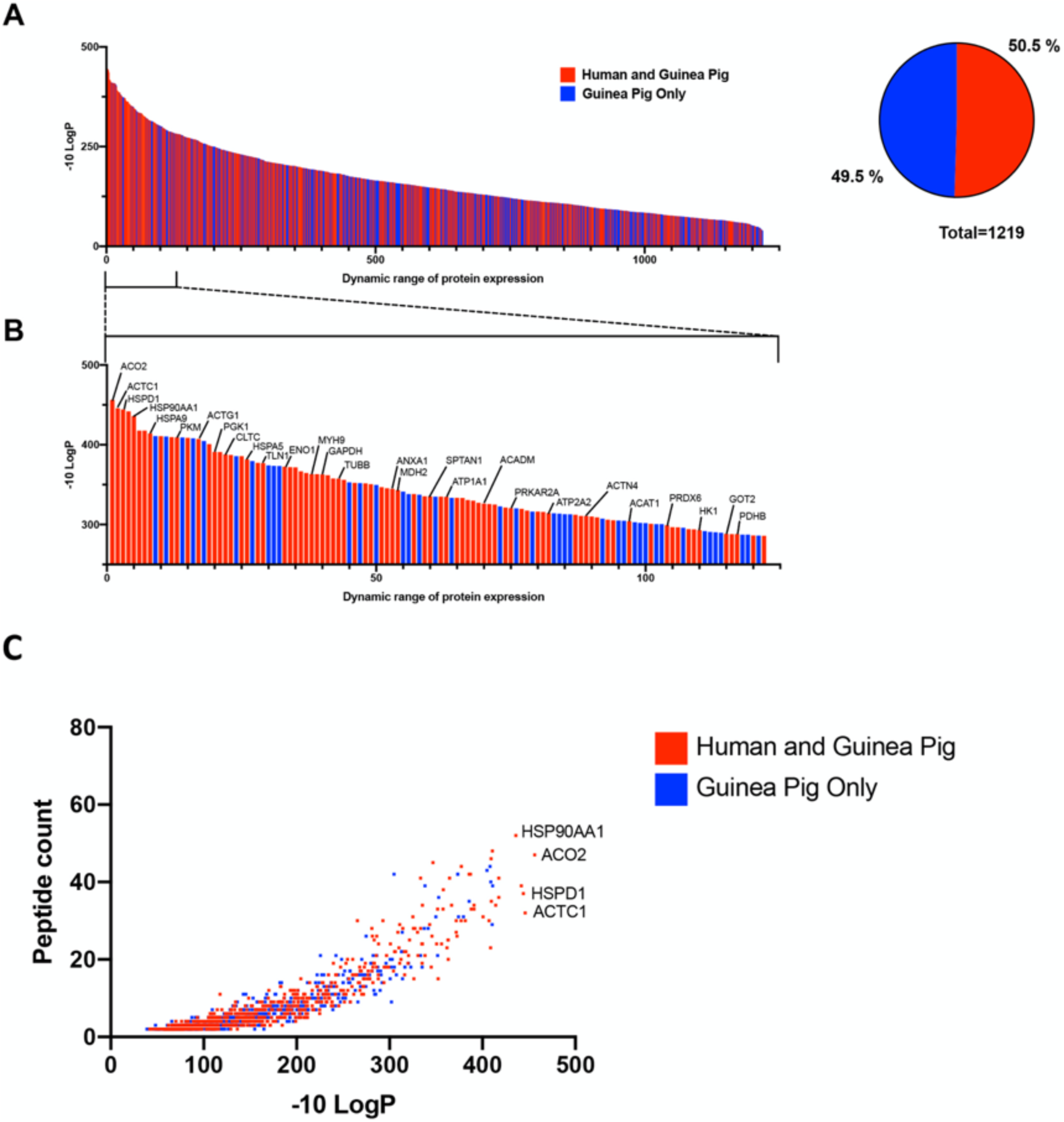
**A**, Dynamic range of peptides identified in guinea pig lysosomal fraction arranged according to −10 LogP values for peptides. Red bars indicate matches with genes confirmed in human lysosomal proteome (data reproduced with permission from Tharkeshwar et al.^83^). Total genes identified = 1219. A complete list of the genes identified in this study is provided in Supplementary Table 1. **B**, expanded view of the top 10% of genes shown in A. Individual gene identifiers have been labelled for reference. **C**, Scatter plot to show relationship between −10 LogP values for all genes identified in guinea pig lysosomal fraction and the total peptide count for each gene. Red boxes indicate matches with genes confirmed in human lysosomal proteome (data reproduced with permission from Tharkeshwar et al.^83^). Labels indicate the most highly expressed genes.

**Supplementary Figure 5.**
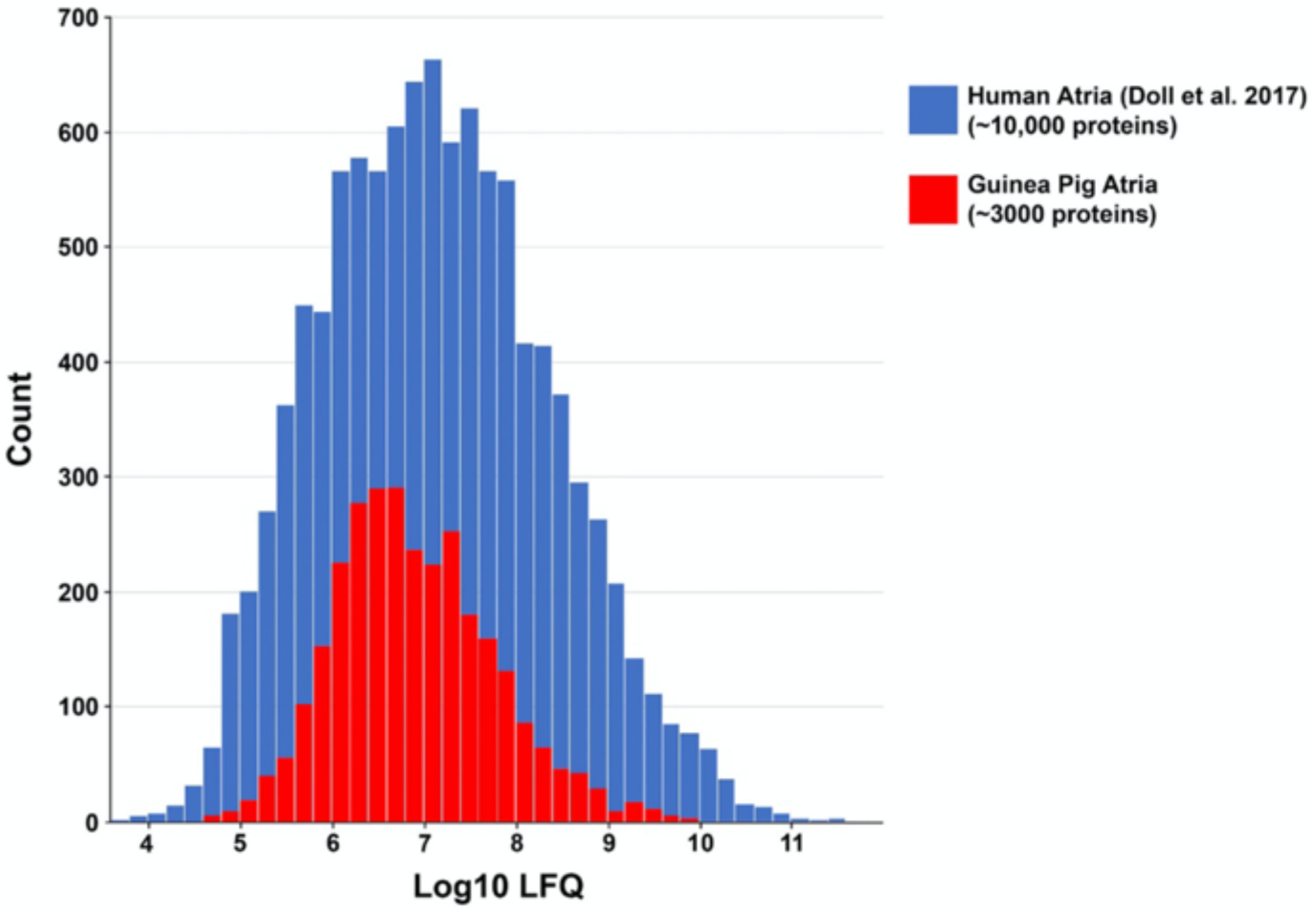
Comparison of proteomic data between human and guinea pig atria. Histograms show the total proteins identified in the TL from our study (red) compared to proteins identified from human left and right atrial tissue samples (blue) in the study by Doll et al.^11^

**Supplementary Figure 6.**
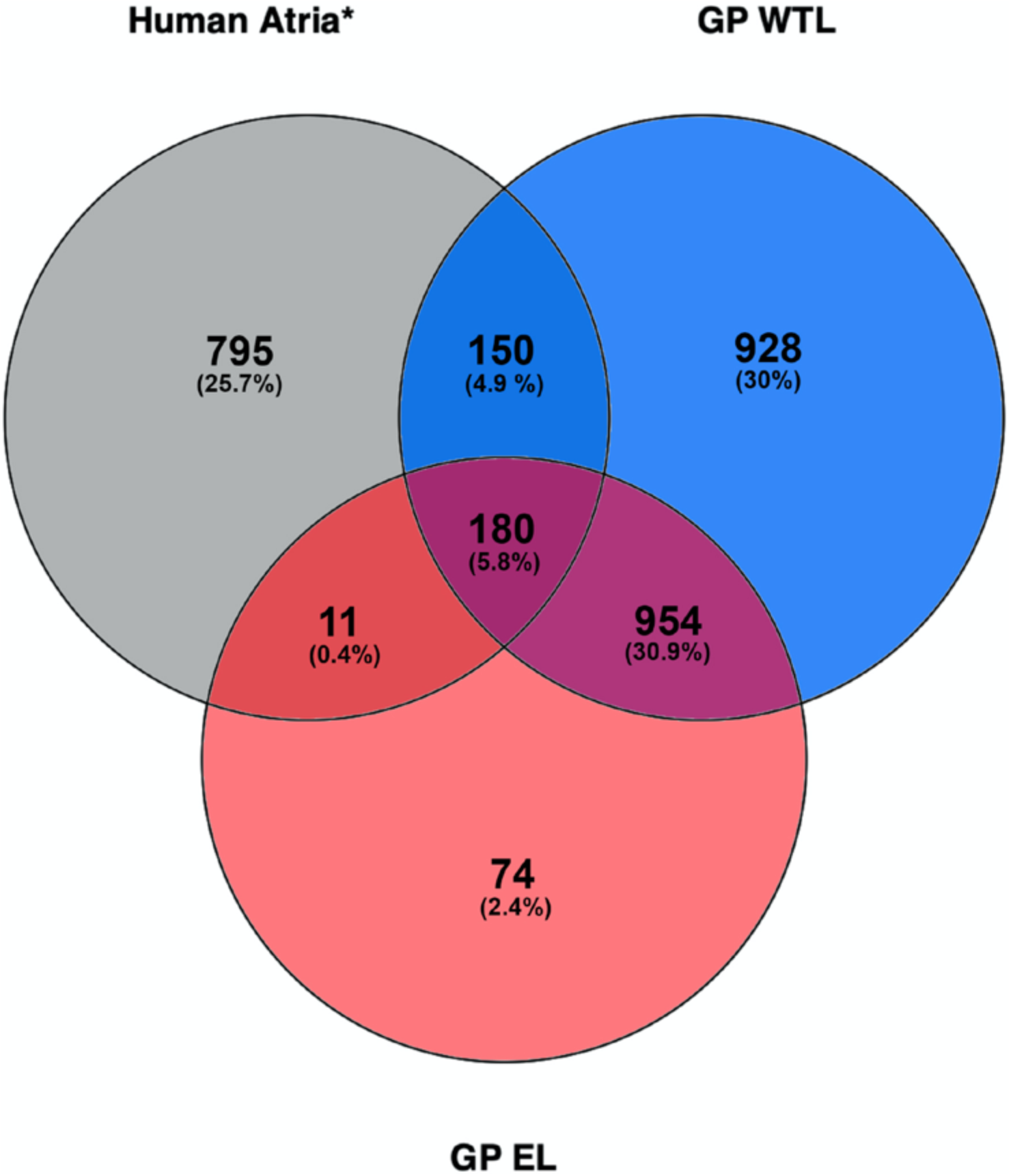
Venn diagram demonstrating overlap in atrial and endolysosomal protein expression in human and guinea pig. Relative overlaps between circles are not shown to scale. GP TL = guinea pig tissue lysate, GP EL = Guinea Pig Endolysosomal Lysate. Data for Human atria based on genes identified by Doll et al^11^ within human atrial tissue samples. Venn diagram produced using Venny 2.1.

